# Different pharmacokinetics of lithium orotate inform why it is more potent, effective, and less toxic than lithium carbonate in a mouse model of mania

**DOI:** 10.1101/2022.05.01.490227

**Authors:** Anthony G. Pacholko, Lane K. Bekar

**Author notes:** **Primary Corresponding Author:** Lane K. Bekar, Ph.D. **Secondary Corresponding Author:** Anthony G. Pacholko, M.Sc.

## Abstract

**Objective:** Lithium carbonate (LiCO) is a mainstay therapeutic for the prevention of mood-episode recurrences in bipolar disorder (BD). Unfortunately, its narrow therapeutic index is associated with complications that may lead to treatment non-compliance. Intriguingly, lithium orotate (LiOr) is suggested to possess uptake properties that would allow for reduced dosing and mitigation of toxicity concerns. We hypothesized that due to differences in pharmacokinetics, LiOr is more potent with reduced adverse effects.

**Methods:** Dose responses were established for LiOr and LiCO in male and female mice using an amphetamine-induced hyperlocomotion (AIH) model; AIH captures manic elements of BD and is sensitive to a dose-dependent lithium blockade. Next, the relative toxicities of LiOr and LiCO were contrasted after 14 consecutive daily administrations.

**Results:** LiCO maintained a partial block of AIH at doses of 15 mg/kg or greater in males and 20 mg/kg or greater in females. In contrast, LiOr elicited a near complete blockade at concentrations of just 1.5 mg/kg in both sexes, indicating improved efficacy and potency. Prior application of an organic-anion transporting polypeptide 1A2 (OATP1A2) inhibitor completely blocked the effects of LiOr on AIH while sparing LiCO, suggesting differences in transport between the two compounds. LiCO, but not LiOr, elicited polydipsia in both sexes, elevated serum creatinine levels in males, and increased serum TSH expression in females.

**Conclusions:** LiOr demonstrates superior efficacy, potency, and tolerability to LiCO in both male and female mice as a result of select transport-mediated uptake.

## Introduction

Lithium salts have been used for more than half a century to combat the psychiatric manifestations of bipolar disorder and, while antipsychotics and anticonvulsants have gained in popularity, lithium remains a frontline therapeutic option (Culpepper, 2014). Of the presently prescribed lithium formulations, lithium carbonate (Li_2_CO_3_; LiCO henceforth) is the most administered, and is one of the most effective medications for the prevention of mood-episode recurrences (Machado-Vieira et al., 2009, Won and Kim, 2017, Malhi et al., 2020, Zivanovic, 2017, Severus et al., 2014, Miura et al., 2014). Unfortunately, LiCO-based therapy displays a narrow therapeutic window with a dose-dependent side effect profile that ranges from mild-to-moderate during short-term use (*e.g*., polydipsia, polyuria) to potentially severe following chronic prescription (*e.g*., nephrogenic diabetes insipidus, hypothyroidism). Consequently, treatment non-adherence is a frequently encountered issue with LiCO therapy (Öhlund et al., 2018).

Lithium orotate (LiC_5_H_3_N_2_O_4_; LiOr henceforth), most notable for its use and advocacy by Hans Nieper in the 1970s (Nieper, 1973), may represent a treatment option that displays lower dosage requirements relative to LiCO with a subsequent reduction in side effect incidence. Nieper proposed that orotic acid was a mineral carrier that could more readily transport inorganic ions – such as lithium, magnesium, or calcium – across biological membranes (Nieper, 1970, Nieper, 1973). Although evidence for enhanced brain availability was initially found (Kling et al., 1978), research into LiOr was discontinued largely due to studies that demonstrated LiOr to increase impairment of kidney function when used at concentrations equivalent to LiCO (Smith and Schou, 1979). While renal toxicity is a concern, we propose that the purported improved bioavailability enables reduced dosage requirements that will mitigate safety concerns. Thus, additional research into the pharmacological properties of LiOr are warranted.

The present study explored the efficacy and potency of LiOr relative to LiCO across a range of concentrations, with the typical therapeutic dose of lithium (adjusted for a murine model) serving as the upper bound. To this end, amphetamine-induced hyperlocomotion (AIH), which has been shown to be attenuated by lithium in a dose-dependent manner (Gould et al., 2007), was used to assess dose requirements of the different lithium compounds.

## Methods and Materials

### Animals

Male and female C57Bl/6NCrl mice (Charles River, Canada) aged 8 weeks were used for all studies. Mice were housed in pairs and kept on a 12-hr light/dark cycle. All experiments were approved by the University of Saskatchewan Animal Research Ethics Board and done according to the Canadian Council on Animal Care.

### Drugs

LiCO – purchased as a powder from Sigma-Aldrich (ON, CA) – was dissolved in distilled water before adjusting the sodium chloride (NaCl) concentration to 0.9%. LiOr was synthesized by combining lithium hydroxide (Sigma-Aldrich; ON, CA), and orotic acid (Sigma-Aldrich; ON, CA) in a 1:1 molar ratio in distilled water; the NaCl concentration was adjusted to 0.9%. For all studies, lithium compound weights are reported as elemental lithium (Li^+^). Dextroamphetamine (*d*A) sulfate tabs (5 mg) were dissolved in saline and administered at 6 mg/kg (0.1 ml/10 g bodyweight). The polyethylene glycol-400 (PEG-400; Sigma-Aldrich; ON, CA) solution was prepared by adding PEG-400 to distilled water in a 1:1 ratio (50% final concentration). PEG-400 was administered 0.1 ml of 50% PEG-400/10 g bodyweight.

### Behavioral tests

#### Amphetamine-induced hyperlocomotion

Similar to humans, mice become hyperactive when administered amphetamine. Mice were administered *d*-amphetamine (*d*A, 6 mg/kg) or saline intraperitoneally (IP), placed into an open field arena (35 × 35 × 35 cm) for 120 minutes, and scored for total locomotion offline using Ethovision XT 11 (Noldus, Wageningen, The Netherlands). For this study, drug efficacy was measured as the ability of the tested lithium compound – which was administered IP 30 minutes prior to application of *d*A – to diminish AIH. For the trials in which PEG-400 is used to block organic anion transporting polypeptides (OATP) (Engel et al., 2012), a 50% PEG-400 solution is delivered via oral gavage 30 minutes prior to the injection of lithium. Locomotion is reported as the percentage of the *d*A response maintained. Locomotion in saline-treated mice represents a full block (17% for males, 13% for females), whereas 100% denotes an absence of blockade (*d*A effect is unimpeded). The Minimal Effective Concentration (MEC) is defined as the lowest lithium concentration used to affect a significant attenuation of AIH.

#### Rotarod (locomotor function)

Male mice were injected with saline or LiCO/LiOr 60 minutes prior to being placed on the rotarod. The rod was accelerated from 4 rpm to 45 rpm over 2 minutes. Each animal was subjected to 4 consecutive trials with the average time to fall of the last three trials recorded for each animal.

#### Forced swim Test

Male mice were injected with saline or LiCO/LiOr 60 minutes prior to being placed into a 4 L beaker filled with 3 L of room temperature water. Activity level was recorded for 8 minutes and analyzed for time spent immobile offline using Ethovision XT 11 software (Noldus, Wageningen, The Netherlands).

### Biochemistry

Animals were anaesthetized using urethane (0.2 mg/ml) and xylazine (150 mg/ml) prior to sacrifice (0.1 ml/10 g body weight). Whole blood and brains were subsequently harvested. Blood was collected via cardiac puncture, deposited into a 1.5 ml Eppendorf, and allowed to clot on ice for 24 hours at 4°C prior to centrifugation at 1500 rcf at 4°C for 15 minutes. Serum aliquots were held at -80°C. Mouse brains were rapidly removed, flash frozen in isopentane, and stored at - 80°C. Frozen brains were ground into a fine powder, mixed with chilled 0.1M PBS + 0.5% tween-20 (5 µg tissue/ml), mechanically homogenized via sonication with three separate 10 second pulses and centrifuged at 20,000 rcf at 4°C for 15 minutes. Supernatants were additionally ultracentrifuged at 200,000 rcf at 4°C for 30 minutes; final supernatants were stored at -80°C.

### Lithium colorimetric assay

Brain and serum Li^+^ content was assayed using a commercially available colorimetric assay (Abcam, item no. Ab235613). Whereas typical clinical laboratories can measure serum Li^+^ down to ∼ 0.3 mM, this colorimetric assay is able to measure Li^+^ content as low at 0.1 mM. Brain samples required adjustment of the sample:sodium-masking-agent:assay-buffer ratio to 15 µl : 15µl : 120 µl from the kit recommended 5 µl : 15µl : 130 µl for serum.

### BUN colorimetric assay

5 µL of serum was diluted 1:9 in 45 µL of distilled water. The diluted samples were assessed for BUN content using a commercially available BUN colorimetric assay (Invitrogen, item no. EIABUN).

### Creatinine ELISA

15 µL of serum was assayed for creatinine content using a commercially available creatinine kinetic colorimetric assay (Cayman chemical, item no. 700460).

### TSH ELISA

30 µL of serum was diluted 1:3 in 90 µL of assay diluent (provided with kit). The diluted samples were assessed for TSH content using a commercially available mouse TSH ELISA kit (Elabscience, item no. E-EL-M1153).

### AST ELISA

2 µL of serum was diluted 1:99 in 200 µL of assay diluent (provided with kit). The diluted samples were assessed for AST content using a commercially available mouse AST ELISA kit (Abcam, item no. ab263882).

### GSK3β activity assay

5 µL of 2 mM LiOr or LiCO were contrasted for their ability to blunt GSK3β *in vitro* activity using a commercially available GSK3β activity-based kinetic colorimetric assay (BPS Bioscience, item no. 79700). The assay required use of the Kinase-Glo Max Luminescent reagent (Promega, item no. V6071).

### Resistivity assay

Resistivity was measured using a patch clamp amplifier (Axopatch 700B; Molecular Devices) and pClamp 10 software (Molecular Devices). A 10-mV voltage jump in current clamp mode was performed during solution transitions from 20 mM LiCl to 20 mM LiOr. The experiment was repeated multiple times with fresh solutions and new glass electrodes.

### Statistics

Data are expressed as mean ± SEM and compared using either one-way or two-way ANOVA with Dunnett’s (one-way) or Bonferroni’s (two-way) post-hoc tests to assess differences between treatment groups (GraphPad Prism V8.1.2; GraphPad Software, Inc. SD, CA). p < 0.05 used as the threshold for significance. SigmaStat 4.0 (Systat Software, Inc. SJ, CA) was used for the construction of the dose-response curves.

## Results

### Amphetamine-induced hyperlocomotion (AIH)

In rodents, the administration of *d*A produces an elevation in central dopamine levels leading to hyperlocomotor activity that reflects manic aspects of bipolar disorder (Sharma et al., 2016, Ashok et al., 2017). Effects of *d*A are known to be sensitive to a dose-dependent lithium block, establishing it as a suitable model for screening mood stabilizers (Gould et al., 2007). The administration of 6 mg/kg *d*A to mice consistently resulted in two distinct peaks of easily quantified hyperlocomotor activity between minutes 5-35 and 70-120 (Fig. 1). The 70-120-minute window was selected to contrast the effects of LiCO and LiOr in these studies because locomotion was particularly robust during this period and potential confounds (IP injection stress, acclimation to novel environment, different pharmacokinetics) for the other peak were of less concern.

**Figure 1:**
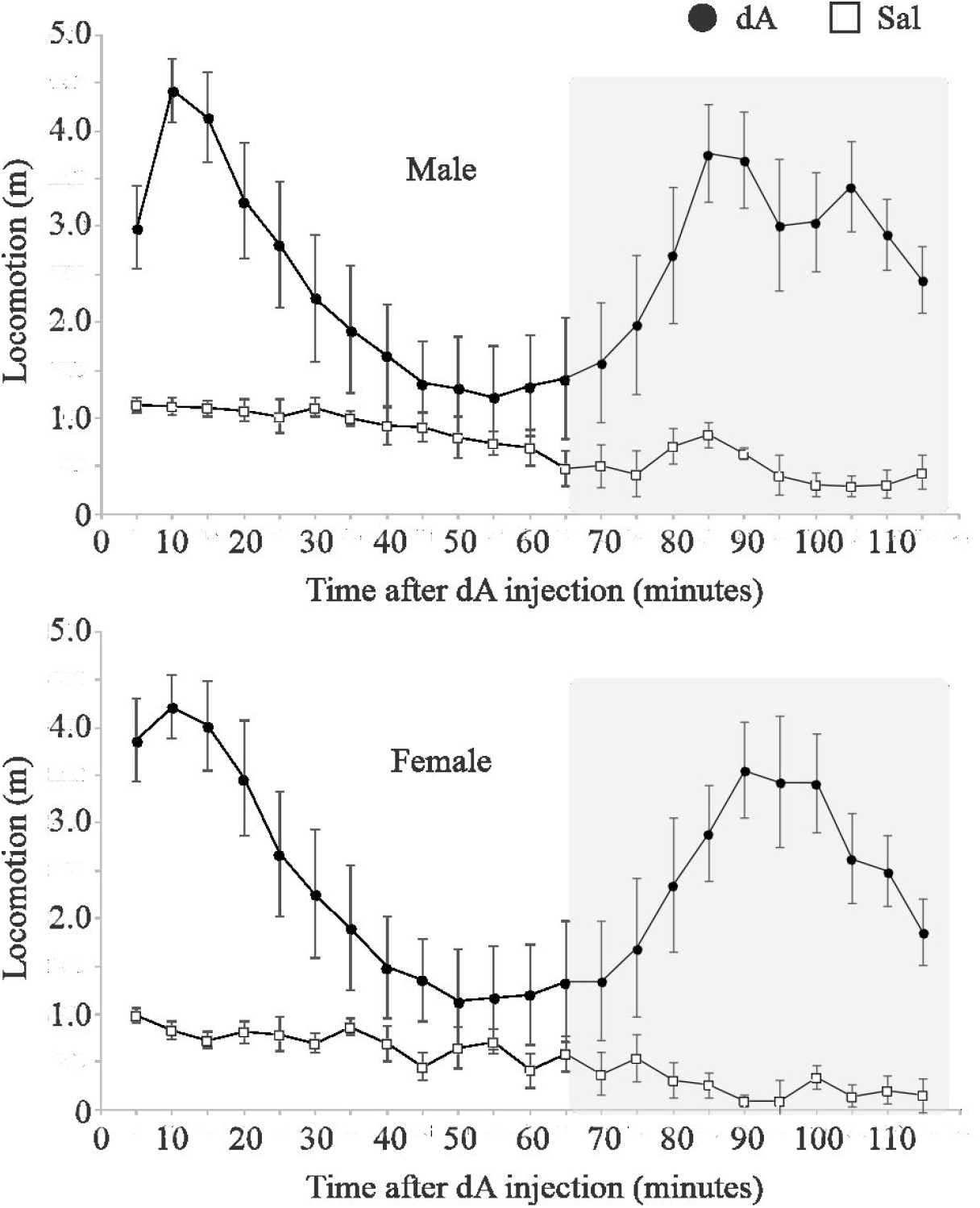
Effects of *d*-amphetamine on locomotor activity. The administration of 6.0 mg/kg *d*A consistently resulted in two periods of hyperlocomotion in both males and females. As the lithium compounds used in this study demonstrated little to no effect within the 1st peak, likely related to timing of injections, the 70-120 minute period was used to contrast the efficacy and potency of each compound. Error bars represent mean ± SEM. N = 5-11. *d*A - d-amphetamine; Sal - 0.9% saline.

### LiOr is more efficacious, potent, and long-lasting than LiCO in the blockade of AIH

To assess the ability of LiCO/LiOr to attenuate AIH, we injected the compounds at various concentrations 30 minutes prior to administration of *d*A. We found that single administration of either LiCO or LiOr blunted AIH in a dose-dependent manner from minutes 70-120 post-*d*A, with LiOr demonstrating a substantially reduced minimal effective concentration (MEC) relative to LiCO in both males and females (Fig. 2A). In males, the MEC was 15 mg/kg for LiCO, and 1.5 mg/kg for LiOr (Fig. 2A, top). Interestingly, the MEC of LiCO demonstrated a rightward shift from 15 mg/kg to 20 mg/kg in females while the MEC for LiOr remained at 1.5 mg/kg (Fig. 2A, bottom). Concurrent with these reduced dose requirements, the strength of blockade elicited by LiOr (75.46 ± 16.95% males; 97.45 ± 6.78% females) was considerably greater than that produced by LiCO (67.18 ± 9.64% males; 82.4 ± 3.99% females), especially in the females. In fact, the MEC for LiOr (1.5 mg/kg in both males and females) elicited a more robust block than the maximum dose for LiCO.

**Figure 2:**
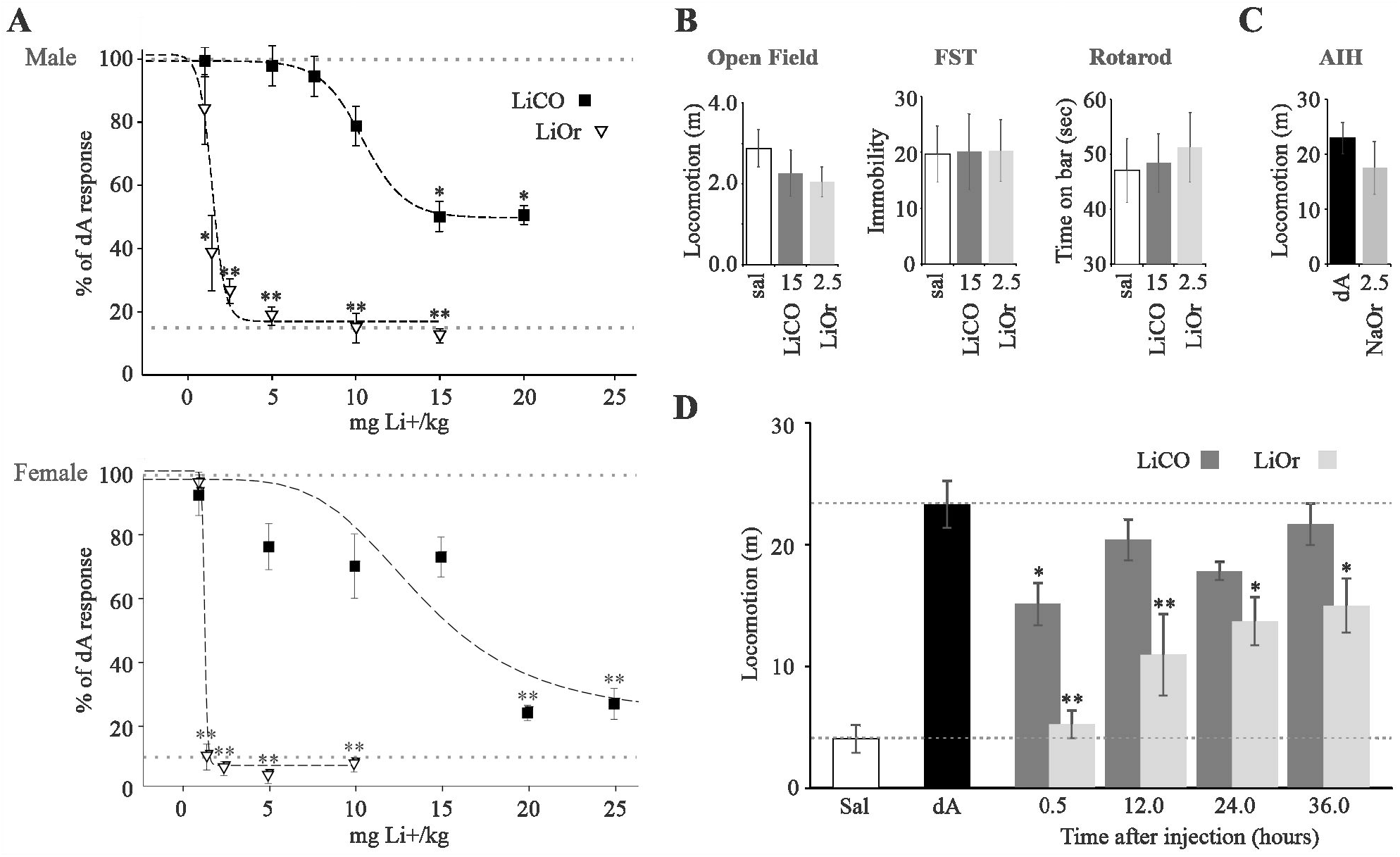
Comparison of LiOr and LiCO effects on lithium-sensitive amphetamine-induced hyperlocomotion. **A)** LiOr and LiCO were administered 30 minutes before *d*A (6 mg/kg). Locomotor scores were tallied between minutes 70-120. LiCO displayed a minimal effective concentration (MEC) of 15 mg/kg in males, and 20 mg/kg in females. LiOr displayed both greater efficacy and potency than LiCO, as evidenced by a more pronounced blockade of hyperlocomotion, as well as a substantially reduced MEC, regardless of sex (n = 5-11). Dashed lines represent 0% (top; *d*A) and 100% (bottom, near the 13% mark on the y-axis; saline) blockade of hyperlocomotion. **B)** Neither LiOr nor LiCO affected baseline activity or motor capacity in the Open Field Test, Forced Swim Test, or rotarod (n = 4-7). **C)** Sodium orotate (2.5 mg/kg) did not affect the AIH ruling out direct effects of orotate. **D)** A single dose of LiOr (2.5 mg/kg), but not LiCO (15 mg/kg), is able to blunt AIH for over 36 hours post administration (n = 5-6). All lithium concentrations are presented as mg of elemental lithium per kg of body weight. Error bars represent the mean ± SEM. All groups were compared to the *d*A control via one-way ANOVA with Dunnett’s post-hoc testing. *P < 0.05. **P<0.01. LiOr - lithium orotate; LiCO - lithium carbonate; *d*A - d-amphetamine; Sal - 0.9% saline; NaOr - sodium orotate.

It is important to note that no notable changes to baseline activity in the open field and/or impairments to locomotor function in the forced swim and rotarod tests were induced by either LiCO or LiOr in the absence of *d*A in male mice (Fig. 2B), thereby signifying that the suppression of hyperlocomotion induced by the MEC of each compound was not due to any lithium-induced reductions in baseline locomotor activity. Also of note, the improved effects of LiOr relative to LiCO are not attributable to orotic acid alone; sodium orotate had no effect on AIH (Fig. 2C).

Given that LiOr has previously been demonstrated to lead to a progressive increase in central Li^+^ levels over the span of 24-hours, even in the face of falling serum concentrations (Kling et al., 1978), we sought to determine whether LiOr could blunt hyperlocomotion when dosed 12, 24 or 36 hours prior to challenge with *d*A. We observed that 15 mg/kg LiCO failed to elicit a significant effect at any time-point beyond 30 minutes (Fig. 2D). In contrast, 2.5 mg/kg LiOr was found to block 66%, 56%, and 52% of AIH at the 12-, 24-, and 36-hour post *d*A-injection time-points, respectively (Fig. 2C). Thus, LiOr demonstrates improved potency (improved effect at reduced concentrations), efficacy (greater blockade of hyperlocomotion), and duration in the attenuation of hyperlocomotion relative to LiCO.

### Evidence for an altered biodistribution of LiOr relative to LiCO

One of the ways in which LiOr is proposed to differ from LiCO is in its lack of dissociation within physiological solutions. Low resistivity is indicative of a solution that allows current/charge flow. Thus, resistivity can be used to assess the degree of a compound’s dissociation/ionization (the lower the electrical resistivity, the greater the degree of ionization). When the dissociation of LiOr and LiCO was contrasted in distilled water, we found that the electrical resistivity of a 20 mM LiCO solution was markedly lower than that of a 20 mM LiOr solution, indicating that LiCO undergoes a greater degree of ionization (Fig. 3A). These results were confirmed using a GSK3β activity assay, where 2 mM LiCO, but not 2 mM LiOr, resulted in an ∼50% reduction in GSK3β activity (Fig. 3B). As Li^+^ must first be liberated from its carrier to inhibit GSK3β, the lack of inhibition elicited by 2 mM LiOr heavily suggests that the compound did not dissociate into its constituent ions. Of note, the IC50 for lithium-induced inhibition of GSK3β *in vitro* is ∼2 mM (Kirshenboim et al., 2004).

**Figure 3:**
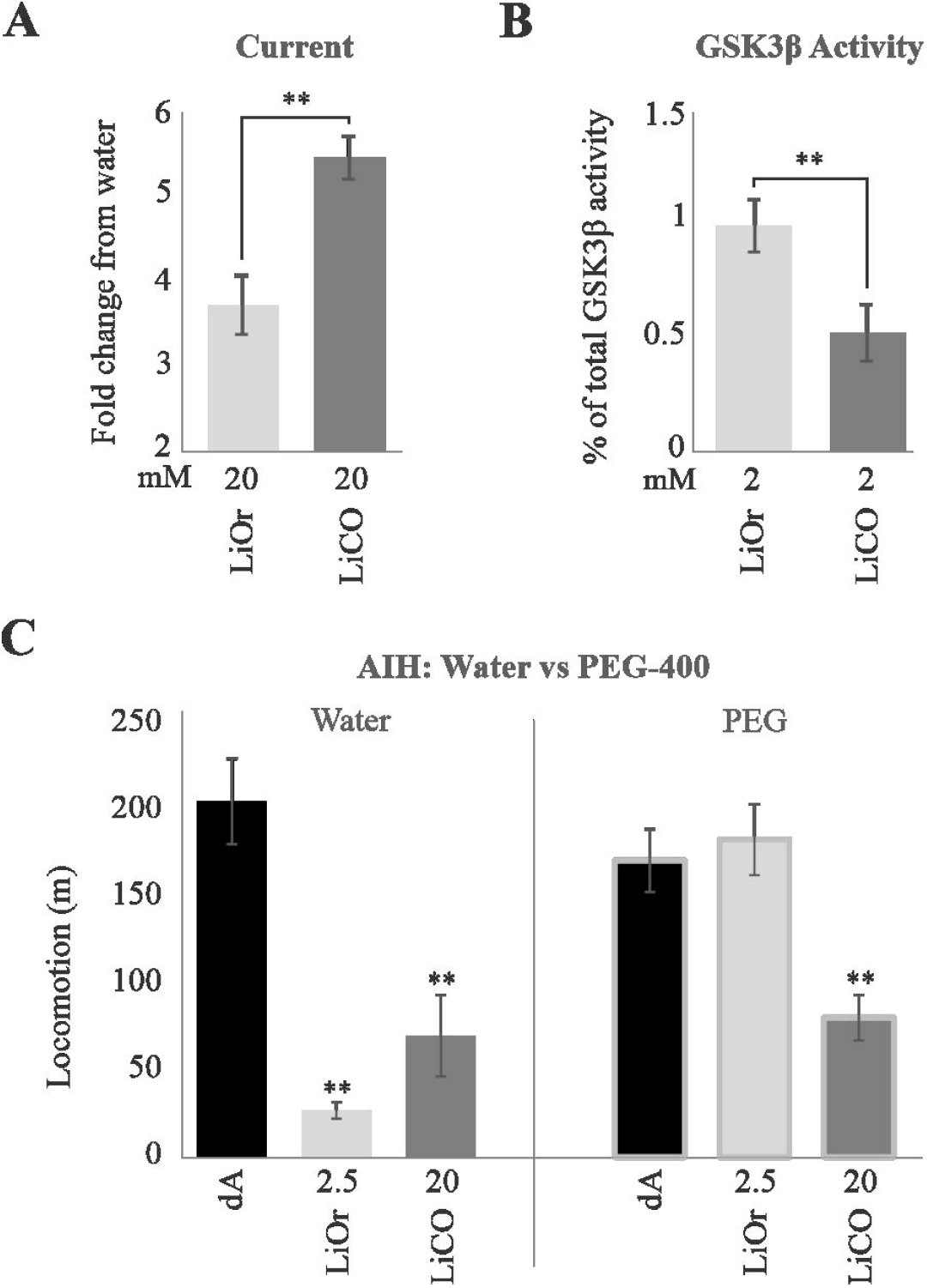
Comparison of LiOr and LiCO differences in biodistribution. **A)** The application of small voltage steps to mixtures containing either 20 mM LiCO or LiOr dissolved in water revealed a substantially lesser degree of resistivity in the LiCO-containing solution than in the LiOr-containing solution. **B)** 2 mM LiCO, but not LiOr, blunted GSK3β activity by ∼50%, which mirrors the IC50 for lithium-induced inhibition of GSK3β *in vitro*. **C)** Pre-application of PEG-400 30 minutes prior to IP injection of LiOr/LiCO completely obstructed the effects of LiOr (2.5 mg/kg) on AIH, whereas LiCO was unaffected (20 mg/kg; hyperlocomotion was blocked as usual). Error bars represent mean ± SEM. For panels **A** and **B**, LiOr and LiCO were contrasted using an unpaired t-test. Each experiment was run at least twice. *P<0.05, **p<0.01. For panel **C**, all groups were contrasted to the *d*A control via one-way ANOVA with Dunnett’s post-hoc testing. Two-way ANOVA was performed for analysis of interactions. n=4-5/group. LiOr – lithium orotate; LiCO – lithium carbonate; AIH – amphetamine-induced hyperlocomotion.

If LiOr does not readily dissociate into orotic acid and Li^+^, then it likely moves throughout the body in a different manner than LiCO (becomes Li^+^ and CO_3_^2-^). Organic anion transporting polypeptides (OATPs) may be involved in transport of LiOr owing to their affinity for large hydrophobic organic anions and abundant localization within both the brain and blood-brain-barrier (BBB) (Roth et al., 2012). Intriguingly, some OATPs, such as OATP1A2 (Oatp1a1 and Oatp1a4 in mice), have been shown to interact with neutral molecules in addition to anions (Schäfer et al., 2021). To explore the potential role of OATPs in the uptake and subsequent efficacy of LiOr, we probed the ability of PEG-400 – a specific inhibitor of OATP1A2/Oatp1a1/Oatp1a4 (Engel et al., 2012) – to affect the efficacy of LiOr and/or LiCO in the attenuation of AIH. The application of 50% PEG-400 via oral gavage 30 minutes prior to IP injection of lithium completely prevented the blockade of AIH ordinarily induced by administration of 2.5 mg/kg LiOr, whereas the 20 mg/kg dose of LiCO was unaffected and continued to blunt AIH as expected (Fig. 3C; two-way ANOVA *interaction*, p = 0.0002).

### In contrast to LiCO, LiOr shows no effect on water intake or kidney and thyroid function

The potential early adverse effects of LiOr and LiCO on kidney and thyroid health – characterized, in part, by aberrant serum TSH, AST, BUN and/or creatinine levels – were contrasted in male and female mice at concentrations of 1x, 2x or 3x the MEC (MEC was 1.5 mg/kg for LiOr and 15 mg/kg for LiCO) once daily for 14 consecutive days. When allometrically scaled, the LiCO concentrations used herein roughly correlate to the therapeutic range employed in humans, *i.e*., 15-45 mg Li^+^/kg in mice translates to ∼400-1200 mgs of total LiCO in an adult patient (correction factor ratio for human to mouse scaling)(Nair and Jacob, 2016). All mice were sacrificed on the 15^th^ day, 24 hours after receiving their final lithium dose.

As polydipsia is a frequent adverse effect of lithium use, we compared the water intake of mice treated with either compound. LiCO, but not LiOr, elicited polydipsia when administered at concentrations greater than or equal to 2x the MEC, with the first signs of excessive water intake observed on day 5 for the 3x dose and day 10 for the 2x dose (Fig. 4A). While the dose-dependent effects of LiCO were similar in each sex, the degree of induced polydipsia was more pronounced in males and demonstrated a progressive increase over time at all concentrations (Fig. 4A, top), whereas water intake plateaued on days 10-through-15 in females treated with the 3x dose (Fig. 4A, bottom). Body weight was unaffected by either treatment (Fig. 4B).

**Figure 4:**
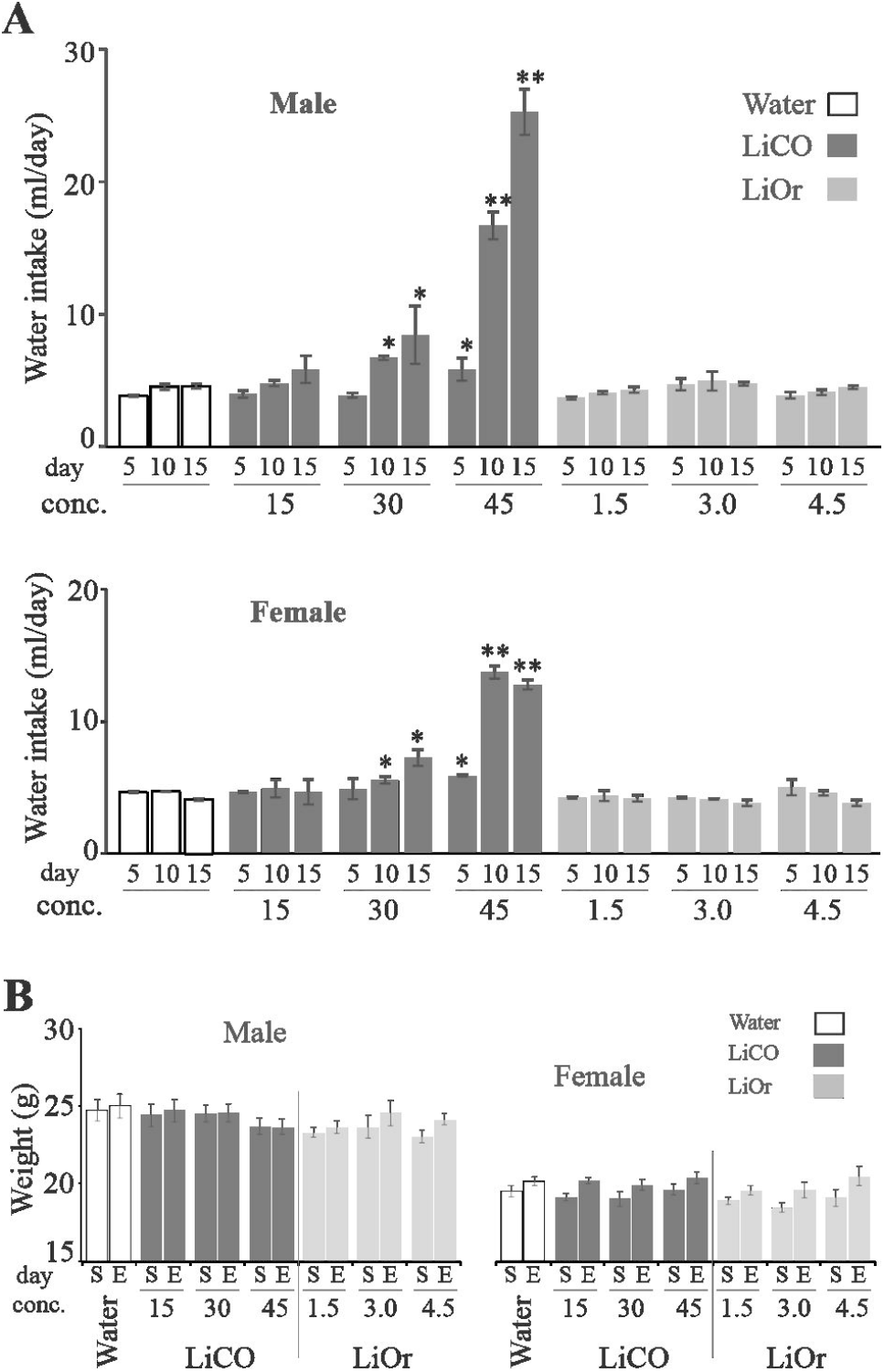
Comparison of LiCO and LiOr effects on water intake in male and female mice. **A)** All compounds were delivered via oral gavage once daily for 15 consecutive days. Water intake was measured every 5 days. LiCO elevates water intake in a concentration-dependent manner. No effects of LiOr on water intake were noted. The polydipsia induced by LiCO at 3x MEC (45 mg/kg) was more substantial in males than in females, with the females demonstrating a plateau at the 3x MEC concentration on days 10-15. **B)** End weight did not significantly differ from start weight in any of the groups within either cohort. Error bars represent the mean ± SEM. For the assessment of water intake **(A)**, all groups were compared to the water-sustained control group via one-way ANOVA with Dunnett’s post-hoc testing. Weight gain or loss was assessed via one-way ANOVA with Tukey’s post-hoc testing (starting weights were compared to end weights for each group). *P < 0.05. **P<0.01. n = 6-7/group. LiOr - lithium orotate; LiCO - lithium carbonate; W - water; S - start weight; E - end weight.

Next, we assessed treatment effects on serum BUN and creatinine, which are waste products used to assess kidney function. Consistent with the lack of effect on polydipsia, LiOr did not alter serum creatinine levels, even when employed at concentrations three-fold greater than its MEC (Fig. 5A, right). In contrast, we observed that the 3x dose of LiCO significantly elevated creatinine levels above control in the male cohort (Fig. 5A, left). Serum BUN levels were unaffected (Fig. 5B).

**Figure 5:**
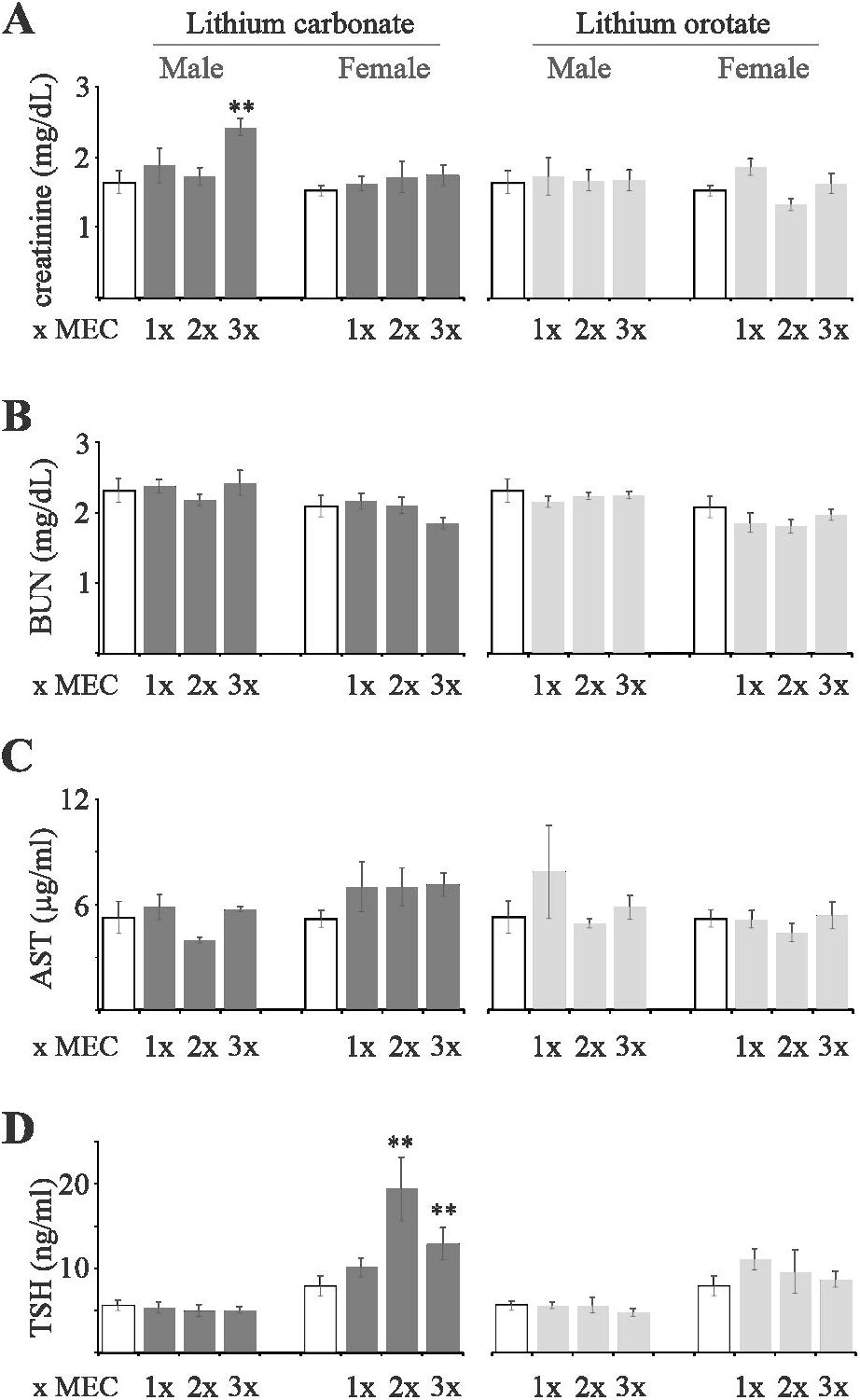
Comparison of LiCO and LiOr effects on traditional markers of thyroid and renal health/function in male and female wild-type mice. **A)** LiCO elevated creatinine levels in males, but not females, when dosed at 3x the MEC (45 mg/kg); LiOr had no effect. **B)** No effects of serum BUN were noted as result of treatment with either LiOr or LiCO. **C)** Although AST was not significantly altered by any of the LiCO concentrations, trends toward significance were noted in the female cohort; males were not affected. **D)** TSH levels were elevated in females, but not males, at all concentrations of 30 mg/kg or greater. All compounds were delivered via oral gavage once daily for 15 consecutive days. Concentrations are based on 1x, 2x or 3x the MEC for each compound, as determined during the SI protocol; thus, concentrations of 1.5, 3 and 4.5 mg/kg were used for LiOr (1.5 mg/kg = MEC for LiOr), while concentrations of 15, 30 and 45 mg/kg were employed for LiCO. When allometric scaling is considered, 15 mg/kg and 45 mg/kg of LiCO (represented as elemental lithium/kg) roughly correlates to the lower and upper bounds of the compound’s therapeutic window. Error bars represent the mean ± SEM. Individual treatment effects, as well as any interaction between drug and sex, were assessed through two-way ANOVA. All male and female LiCO and LiOr groups were compared to their respective water control via Bonferroni post-hoc testing. *P < 0.05. ***P<0.01. n = 3-6/group for AST, 5-7/group for TSH, and 6- 7/group for BUN and creatinine. BUN - blood-urea-nitrogen; TSH - thyroid stimulating hormone; AST - aspartate aminotransferase; LiOr - lithium orotate; LiCO - lithium carbonate.

No alterations in serum AST content, which can indicate kidney and/or liver damage when increased, were observed for either LiCO or LiOr (Fig. 5C).

Finally, we assessed the impacts of each lithium treatment on serum TSH, which serves as a clinical marker of lithium-induced hypothyroidism. LiCO, but not LiOr, elevated TSH expression in females, whereas the male mice were unaffected (Fig. 5D). The differences between the male and female cohorts were associated with an interaction between sex and treatment (two-way ANOVA *interaction*, p<0.001), which suggests that the effects of LiCO on TSH are sex-dependent.

In summary, LiOr did not elicit any adverse effects on either water intake or serum biomarkers of lithium toxicity, even when dosed at three times its MEC. In contrast, treatment with LiCO elevated serum TSH in females, serum creatinine in males, and polydipsia in both males and females.

### LiOr retains efficacy at concentrations that are undetectable within the serum

We observed that serum Li^+^ levels were elevated in mice treated with LiCO or LiOr relative to saline alone in both males and females (Fig. 6A; saline not shown due to failure to meet the 0.1 mM detection threshold). Interestingly, the serum Li^+^ levels resultant of LiOr administration were only detectable at concentrations 1.67 (males) or 3.33 (females) times greater than the MEC of 1.5 mg/kg, whereas LiCO generated detectable levels when employed at concentrations well below its MEC of 15 mg/kg (Fig. 6A).

**Figure 6:**
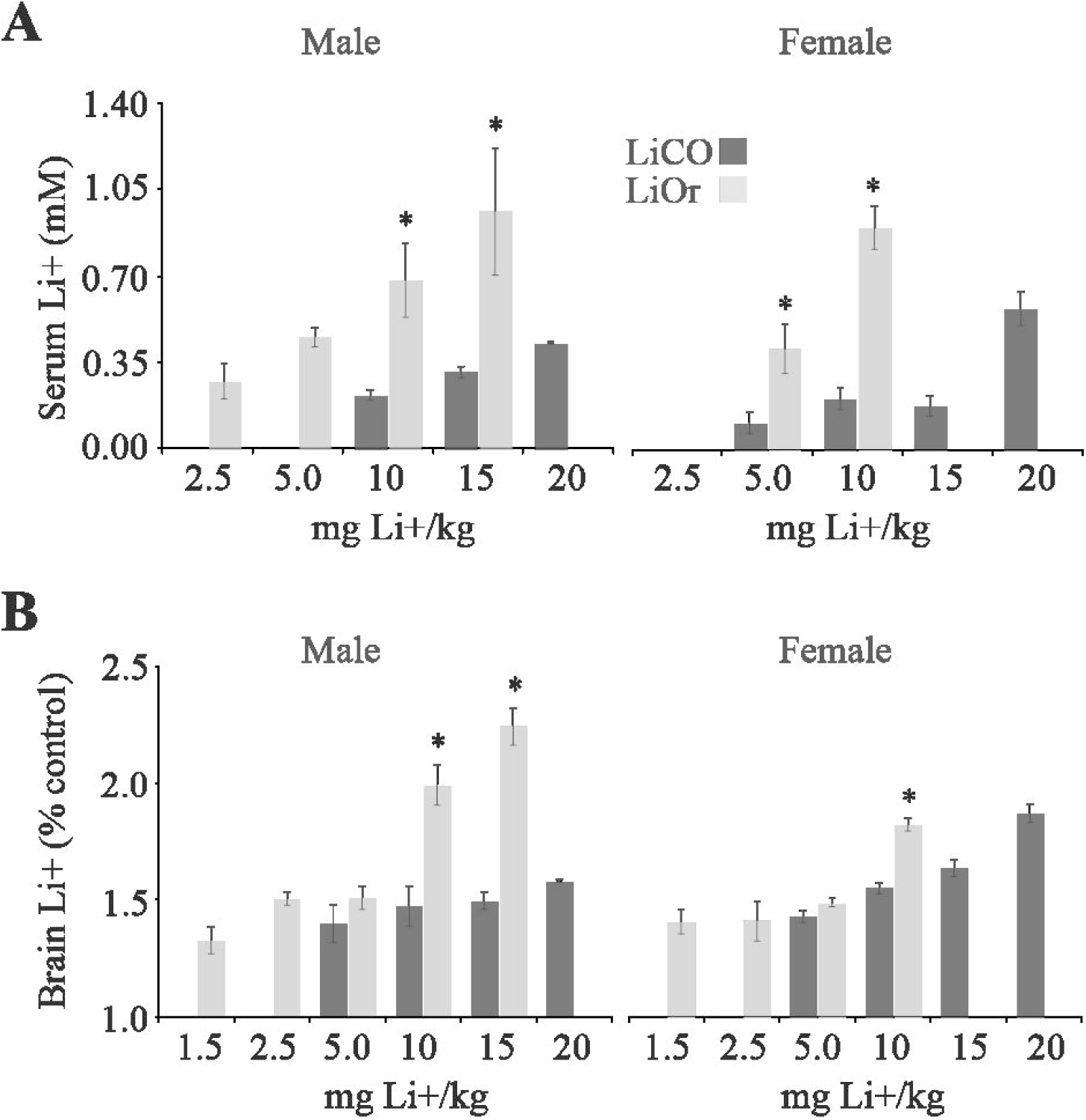
LiOr yields greater brain and serum lithium levels than LiCO. **A**) LiOr produced higher serum lithium levels at all matching concentrations (5-10 mg/kg), regardless of sex. LiOr failed to reach the detection limit at doses lesser than or equal to 1.5 mg/kg, whereas LiCO fell below the detection limit at 5 mg/kg and 10 mg/kg in females and males, respectively. **B)** Brain lithium levels were elevated in all mice treated with lithium, regardless of compound or sex. LiOr produced higher brain lithium levels than LiCO at administered concentrations of 10 mg/kg or greater in both male and female mice. Error bars represent mean ± SEM. Brain lithium content for all groups was contrasted to the saline control via one-way ANOVA with Dunnett’s post-hoc (matched concentrations, e.g., LiOr 5 versus LiCO 5, were compared using Tukey’s post-hoc test). *P<0.05, **p<0.01. n=4-7/group. LiOr – lithium orotate; LiCO – lithium carbonate. The detection limit was 0.1 mM.

As LiOr has previously been found to increase central Li^+^ levels relative to LiCO (Kling et al., 1978), brain Li^+^ was contrasted in LiCO and LiOr treated mice. We found that both lithium compounds significantly increased brain Li^+^ levels relative to control at all tested concentrations, with LiOr-treated mice displaying higher brain Li^+^ levels than LiCO at concentrations greater than or equal to 10 mg/kg in both males (fig. 6B left) and females (fig. 6B right).

## Discussion

Despite early evidence of reduced dosage requirements relative to LiCO, the use of LiOr in psychiatric applications has gone largely unexplored over the past 50 years (Kling et al., 1978, Nieper, 1973, Sartori, 1986). Using the AIH model of mania (Fig. 1) – which was selected for its high-throughput and established dose-sensitivity to lithium – we found LiOr to be more potent, efficacious, and long-lasting than LiCO in the blockade of hyperlocomotion (Fig. 2). These differences likely relate to the observations that LiOr did not readily dissociate in solution and appeared to utilize alternative transport mechanisms (Fig. 3). Furthermore, we observed an absence of adverse effects on markers of kidney and thyroid health with LiOr (Figs. 4, 5) supporting the notion that at the lower effective dosages, LiOr is a safer alternative to LiCO.

The lines of evidence supporting the translational potential of the improved potency and efficacy of LiOr relative to LiCO are manifold. First, the actions of lithium against hyperlocomotion are dependent upon amelioration of the amphetamine-induced increase in GSK3β signaling downstream of dopamine receptors (Beaulieu et al., 2008); amphetamine elevates GSK3β activity, in part, through inhibition of the dopamine transporter (DAT), which subsequently results in an enhanced dopaminergic tone. As excessive GSK3β output (Jope, 2011, Jope and Roh, 2006, Muneer, 2017, Yu and Greenberg, 2016), increased expression of dopamine receptors, and reduced availability of DAT have been implicated in BD pathogenesis (Anand et al., 2011, Milienne-Petiot et al., 2017, van Enkhuizen et al., 2014, Ashok et al., 2017), the odds that the improved potency/efficacy of LiOr noted in the AIH model will translate to the human condition appear promising. Second, while no clinical trials for the use of LiOr in BD have been conducted, the disparity in the MEC between LiOr and LiCO pertaining to blockade of AIH is mirrored in studies exploring their efficacy in the cessation of alcohol abuse. LiOr has shown success in reducing alcohol consumption when administered daily for 6 months at a dose of 150 mg/day (∼6.4 mg of Li^+^)(Sartori, 1986), whereas LiCO is either mildly efficacious (Fawcett et al., 1987) or outright ineffective (Dorus et al., 1989) when employed at substantially greater doses (>600 mg/day; ∼112 mg of Li^+^). Finally, the MEC for LiCO translates to ∼500+ mg of LiCO/day in an 80 kg man or 70 kg woman when scaled from rodent to human, which aligns with the lower end of the effective range employed during lithium therapy and supports the idea that the dose necessary for blockade of AIH roughly correlates with the therapeutic dosages used for the control of mania.

In concert with its reduced dosage requirements, LiOr did not elicit any adverse kidney health-related outcomes (elevated BUN, creatinine, polydipsia, etc.) at doses up to three-fold greater than its MEC. Given the positive association between serum Li^+^ levels and toxicity (Malhi, 2015), it is possible that this tolerability is attributable to the fact that the MEC for LiOr does not give rise to detectable levels of Li^+^ within the serum. Additionally, the increased duration of effect noted for LiOr (12-36 hours post-administration) may result in a smoother serum Li^+^ curve over time that minimizes the incidence of “Li^+^ spikes”; lithium-induced toxicity is worsened by acute spikes in serum Li^+^ levels (Malhi, 2015). In contrast, LiCO elicited elevations in serum creatinine content concurrent with severe polydipsia in male mice, suggesting an impaired ability to concentrate urine in a manner that may reflect vasopressin resistance, which is a frequent complication of lithium use. While the differences between LiOr and LiCO at this early time point are insufficient to definitively state that LiCO will display toxicity during chronic treatment while LiOr will not (Gitlin, 2016), the failure of LiOr to elicit water-balance-associated side-effects, which are frequently encountered during the early stages of lithium therapy, suggests that LiOr will demonstrate superior long-term tolerability. This supposition is supported by the absence of any reported cases of serious side effects in over 40 years of LiOr use in North America (Devadason, 2018).

In line with our observations pertaining to kidney health, LiCO, but not LiOr, elicits an elevation in serum TSH at therapeutically relevant concentrations in female mice, which suggests that LiOr may spare thyroid output. Thus, the seemingly improved tolerability of LiOr may be of particular benefit to female BD patients, who are known to be at greater risk for the development of lithium-induced hypothyroidism than their like-aged male counterparts (Henry, 2002).

Opposing our submission of improved tolerability, some have suggested the improved efficacy of LiOr to be attributable to reduced glomerular filtration rates that ultimately culminate in worsened renal health outcomes (Smith and Schou, 1979, Kling et al., 1978). However, our present results are supported by a recent 28-day toxicological evaluation of LiOr at doses up to 400 mg/kg/day in rats (elemental Li^+^ ∼ 15 mg/kg/day) in which no adverse effects were found (Murbach et al., 2021). Furthermore, the most well-known case of LiOr-induced toxicity seemingly highlights its safety. In 2007, a case report detailing a scenario in which 18 LiOr tablets (3.83 mg Li^+^/tablet) were ingested showed that the patient merely displayed nausea, minor tremors, and normal vital signs, with all symptoms resolving after 3 hours of observation without intervention (Pauzé D and Brooks D, 2007). The MEC for LiOr in the attenuation of AIH is roughly equivalent to just 2-3 LiOr tablets in a reasonably sized human man (80 kg) or woman (70 kg).

The early proponents of LiOr argued that the improved efficacy of the compound is linked to the utilization of uracil-specific transport systems as well as affinity for tissues highly expressing the pentose phosphate pathway (Nieper, 1970, Nieper, 1973). The structure of LiOr closely resembles that of 5-fluorouracil, which is a non-charged pyrimidine known to be an exogenous substrate for the ubiquitously expressed equilibrative nucleotide transporters (Wohlhueter et al., 1980). While intriguing, the greatest support for the notion that LiCO and LiOr differ in terms of transport may be our findings that a) LiOr does not readily dissociate into its constituent ions, and that b) PEG-400 completely prevents inhibition of AIH by LiOr while sparing the inhibition by LiCO. As PEG-400 and LiOr were administered via different routes (OG and IP, respectively), it is likely that the effects of PEG-400 are chiefly attributable to its inhibition of OATPs. OATP1A2 (Oatp1a1 and Oatp1a4 in mice) appears to be of particular importance, as it is localized within neurons, glial cells, and the endothelium of the BBB (Schäfer et al., 2021), and is a specific target for inhibition by PEG-400 (Engel et al., 2012). Thus, while LiCO requires large serum Li^+^ concentrations in order to “drive” Li^+^ into cells, the putative transport- and dissociation-related properties of the orotic acid carrier may reduce dose requirements by allowing delivery of Li^+^ directly to the intracellular target site, as was originally proposed by Hans Nieper in the early 1970s (Nieper, 1973).

In closing, the reduced dosage requirements observed for LiOr in the present study appear to dispel the concerns regarding renal toxicity raised in 1979 (Smith and Schou, 1979) where they used equal elemental Li+ concentrations of the two forms (CO and Or), as well as ameliorate the dose-dependent, compliance-disrupting side-effects associated with current LiCO therapy. Given the potency, efficacy, apparent tolerability, and wide-spread availability of this over-the-counter nutraceutical, clinical trials for the use of LiOr are warranted.

## Acknowledgments

The present work was supported by a College of Medicine Research Award from the University of Saskatchewan.

